# Real-Time Measurement of Stimulated Dopamine Release in Compartments of the Adult *Drosophila melanogaster* Mushroom Body

**DOI:** 10.1101/2020.06.29.177675

**Authors:** Mimi Shin, Jeffrey M. Copeland, B. Jill Venton

## Abstract

*Drosophila melanogaster*, the fruit fly, is an exquisite model organism to understand neurotransmission. Dopaminergic signaling in the *Drosophila* mushroom body (MB) is involved in olfactory learning and memory, with different compartments controlling aversive learning (corner) vs appetitive learning (medial tip). Here, the goal was to develop techniques to measure endogenous dopamine in compartments of the MB for the first time. We compared three stimulation methods: acetylcholine (natural stimulus), P2X_2_ (chemogenetics), and CsChrimson (optogenetics). Evoked dopamine release was measured with fast-scan cyclic voltammetry in isolated adult *Drosophila* brains. Acetylcholine stimulated the largest dopamine release (0.40 μM), followed by P2X_2_ (0.14 μM), and CsChrimson (0.07 μM). With the larger acetylcholine and P2X_2_ stimulations, there were no regional or sex differences in dopamine release. However, with CsChrimson, dopamine release was significantly higher in the corner than the medial tip, and females had more dopamine than males. Michaelis-Menten modeling of the single-light pulse revealed no significant regional differences in K_m_, but the corner had a significantly lower V_max_ (0.12 μM/s vs. 0.19 μM/s) and higher dopamine release (0.05 μM vs. 0.03 μM). Optogenetic experiments are challenging because CsChrimson is also sensitive to blue light used to activate green fluorescent protein, and thus, light exposure during brain dissection must be minimized. These experiments expand the toolkit for measuring endogenous dopamine release in *Drosophila*, introducing chemogenetic and optogenetic experiments for the first time. With a variety of stimulations, different experiments will help improve our understanding of neurochemical signaling in *Drosophila*.

## Introduction

*Drosophila melanogaster*, the fruit fly, is a model system for the field of neuroscience. The fruit fly’s brain consists of around 100,000 neurons that modulate complex behaviors, such as learning and memory ^1^. Although flies and vertebrates differ greatly in brain anatomical structures, many of basic elements and functions of neurotransmission are well conserved between both organisms ^2^. For example, flies use many of the same neurotransmitters as mammals including dopamine, serotonin, glutamate, acetylcholine, etc. In *Drosophila*, eight clusters of dopaminergic neurons have been identified that are distributed throughout the brain, projecting to different regions, including mushroom body (MB) and central complex ^3^. Three dopaminergic clusters project to the MB lobes which are architecturally compartmentalized into 15 discrete sections depending on the MB output and dopaminergic neuron innervation ^3,4^. Protocerebral anterior medial (PAM) and protocerebral posterior lateral 2 (PPL2) neurons project to the tip of horizontal lobes and calyx, respectively, and PPL1 neurons project to the vertical lobes, heel, and peduncle of MB ^5,6^. Each dopamine projection independently regulates synaptic activity in MB, promoting approach or avoidance behaviors, olfactory valence, and memory strength during olfactory learning ^7,8^. Studies to investigate the dopaminergic system and underlying mechanism of learning and memory heavily rely on genetics, behavior assays, and calcium imaging ^9–11^. These approaches reveal the function of specific genes, but lack the ability to monitor real-time changes of endogenous neurotransmission in *Drosophila* brains.

Electrochemical detection is a good strategy for real-time detection of neurotransmitters because it has high temporal resolution.^12^ Microelectrodes, typically less than 10 μm, are small enough to implant in specific target regions in the *Drosophila* central nervous system ^13^. Amperometry and chronoamperometry were previously used with carbon-fiber microelectrodes (CFMEs) to measure either exogenously applied dopamine or nicotine-stimulated octopamine release in adult fly brains ^14,15^. Although, both amperometry and chronoamperometry can acquire rapid measurements, they suffer from lack of selectivity ^16^. Fast-scan cyclic voltammetry (FSCV) is commonly used to measure neurochemicals *in vivo* and *in vitro* because it offers subsecond temporal resolution and a CV, fingerprint for the measured analyte providing provides selectivity ^17–20^. Our group has previously employed FSCV at CFMEs to monitor ontogenetically-evoked endogenous neurotransmitters in the larval ventral nerve cord (VNC) ^17,21–24^. In adult flies, we measured acetylcholine-evoked dopamine in the central complex of adult brain; however, the MB has not been explored ^24^.

Several stimulation methods have been applied in *Drosophila* to elicit endogenous neurotransmitter release. Optogenetics uses genetic tools, such as Gal4-UAS system, to express light-gated ion channels controlling specific neuronal activity that results in synaptic neurotransmission ^25^. In fly larvae, CsChrimson, a red-light sensitive channel, was previously used with FSCV detection because red light does not cause electrochemical artifacts as bluelight sensitive Channelrhodopsins does ^21^. In a complementary strategy, chemogenetic channels regulate cellular activity using a chemical stimulant to activate specific, chemical-sensitive ion channels ^26^. P2X_2_, an ATP sensitive cation channel, is not endogenously encoded in flies; thus when it is expressed in neurons, applying extracellular ATP also causes synaptic neurotransmission ^27^. Acetylcholine, an excitatory neurotransmitter in insects, activates nicotinic acetylcholine receptors (nAChRs) and causes rapid synaptic neurotransmission. Acetylcholine binds to presynaptically expressed nAChRs on dopaminergic terminals, regulating dopamine release and does not require any genetic manipulation compared to optogenetics and chemogenetics^24,28^.

The goal of this study was to compare endogenous dopamine release in different compartments of the MB in isolated adult fly brains. Acetylcholine-, P2X_2_-, and CsChrimson-mediated dopamine release were compared in different MB compartments of males and females. Acetylcholine stimulated the highest dopamine release, followed by P2X_2_ and CsChrimson. With CsChrimson, dopamine release was significantly higher in the corner than the medial tip and higher in females than males. Modeling Michaelis-Menten uptake demonstrates regional differences in V_max_ and dopamine release. CsChrimson is sensitive not only to red light but also to other, lower wavelengths and thus minimizing the light exposure during brain dissection is key to performing optogenetic experiments. This work develops three stimulation techniques for *Drosophila* and demonstrates regional and sex differences in dopamine signaling in the MB. Thus, there is now an established toolkit of stimulation methods that can be used to understand rapid neurotransmission in the *Drosophila* adult brain,

## Experimental Methods

### *Drosophila Melanogaster* Brain Tissue Preparation

Detailed procedure on preparing fly strains can be found in supporting information. *Drosophila* adult brain tissue was prepared as described in Shin *et al*. ^24^. Briefly, a 5 to 10-day old adult fly was anesthetized by rapid chilling on a chilled Petri dish for 1 min. Then, the brain tissue was isolated in a chilled dissecting buffer and the glial sheath was carefully removed with sharp tweezers. The harvested brain was then transferred to a Petri dish containing room temperature dissecting buffer and then placed at the bottom of the dish anterior side up. For acetylcholine and P2X_2_ experiments, a CFME was placed either at the heel or medial tip of MB using GFP marker as a guidance to locate the target region, and a pipette filled with either acetylcholine or ATP was positioned approximately 10 to 15 μm away from the CFME tip. Brain tissue was allowed to equilibrate 15 min prior to the data collection. For CsChrimson experiments, the CFME was implanted with minimum light and positioned either at the corner and medial tip without GFP marker, and the optic fiber was placed right above the isolated brain tissue. Brain tissue was rested in dark for 30 min prior to the experiment. After experiments were completed, fluorescent light was used to activate GFP marker to confirm the electrode placement.

### Stimulation of Neurotransmitter Release

A Picospritzer III (Parker Hannfin, Fairfield, NJ) was employed to pressure eject acetylcholine or ATP into the brain tissue. Pipettes were calibrated by picospritzing a droplet in oil, and the diameter of the droplet was measured using DS-Qi2 monochrome CMOS camera and NIS-Elements BR imaging software (Nikon Instrument Ins, Melville, NY). For optogenetic stimulation, red-orange light (617 nm) fiber-coupled LED with a 200 μm core optical fiber cable and LED controllers (ThorLabs, Newton, NJ) were used with to deliver light stimulation. Light stimulating parameters, such as frequency, pulse width, and pulse number, were modulated with Transistor-Transistor Logic (TTL) inputs which were generated by electrical pulses controlled with the TarHeel CV software.

### Statistics Analysis and Modeling

All statistics were performed using GraphPad Prism 8 (GraphPad Software, Inc, La Jolla, CA). All data are reported mean ± standard error of the mean. To model the uptake kinetics of single light pulse evoked dopamine release, MATLAB based MMFIT program was used (provided by Charles Nicholson, New York University School of Medicine), which was applied as described in Li *et al*. ^29^. Single pulse dopamine peak concentration curve was used to model uptake kinetics. 100 ms after the maximum peak concentration curve (three data points) was fitted to extract V_max_ and k_m_ using the following Michaelis-Menten equation:

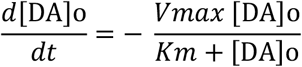

where V_max_ and k_m_ are the Michaelis-Menten uptake constants and [DA]_o_ is the single pulse evoked dopamine peak concentration. This modeling uses a nonlinear curve fitting simplex with a standard root finding algorithm assuming that diffusion does not contribute to dopamine clearance.

## Results and Discussion

### Acetylcholine Mediated Endogenous Dopamine Release in the Mushroom Body of *Drosophila* Brains

The goal of this study was to compare multiple stimulation methods to evoke endogenous dopamine release in different compartments of the mushroom body (MB) in adult *Drosophila* brains. The MB consists of three lobes (Fig. 1A), αβ, α’β’, and γ lobes, and γ lobe is commonly studied to investigate the distinct roles of neuronal circuitry during olfactory learning and memory [39]. Dopamine measurements were made either at the corner, from the γ1 and γ2 compartments, or at the medial tip from the γ4 and γ5 compartments. Dopaminergic neurons were visualized in the MB by crossing dopamine transporter (DAT) driver (DAT-Gal4) flies with a green fluorescent protein responder (UAS-GFP) line to express GFP in neurons containing DAT (Fig. 1B) ^30^. For experiments, a carbon-fiber microelectrode (CFME) and a pipette were placed 10 to 15 μm apart, either in the corner (Fig. 1C) or medial tip (Fig. 1D).

**Figure 1.**
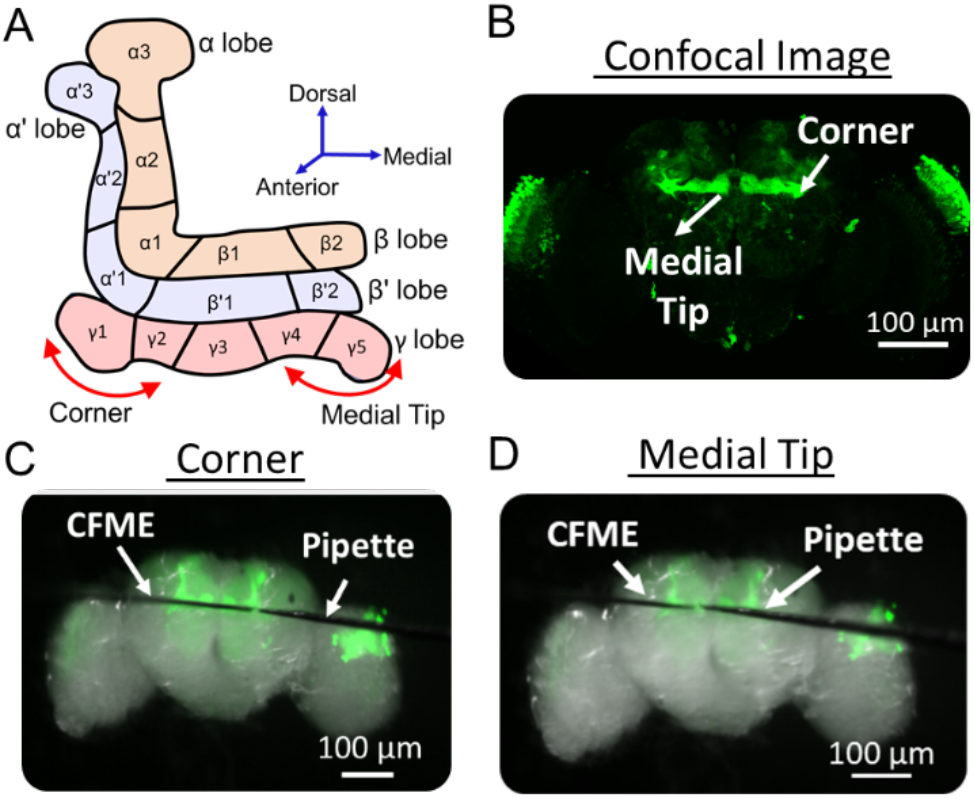
Mushroom bodies in adult *Drosophila melanogaster* brain. (A) A schematic diagram of the mushroom body (MB) with each lobe and compartment labeled. (B) A confocal image of adult brain of a DAT-Gal4, UAS-GFP fly. MB strongly expresses GFP. A microscope image of adult fly brain with a carbon-fiber microelectrode (CFME) and pipette filled with 5 mM acetylcholine placed at the (C) corner or (D) medial tip of MB.

The first stimulation method was acetylcholine, which was pressure injected to stimulate dopamine release in isolated adult *Drosophila* brains. Figures 2A and 2B show representative FSCV data of 1 pmol acetylcholine-evoked dopamine in the corner and the medial tip of MB, respectively. FSCV provides a color plot that shows the voltage, current, and time data in 3 dimensions; dopamine is visualized as the green circle, representing oxidation, and a blue circle, representing reduction (Fig. 2A, bottom). In addition, a current vs. time trace extracted at the oxidation potential shows changes over time (Fig. 2A, top) and the cyclic voltammogram (CV) confirms dopamine is detected (Fig. 2A, insert). Dopamine rapidly increased after acetylcholine was applied, but took about 20 s to return to baseline.

**Figure 2.**
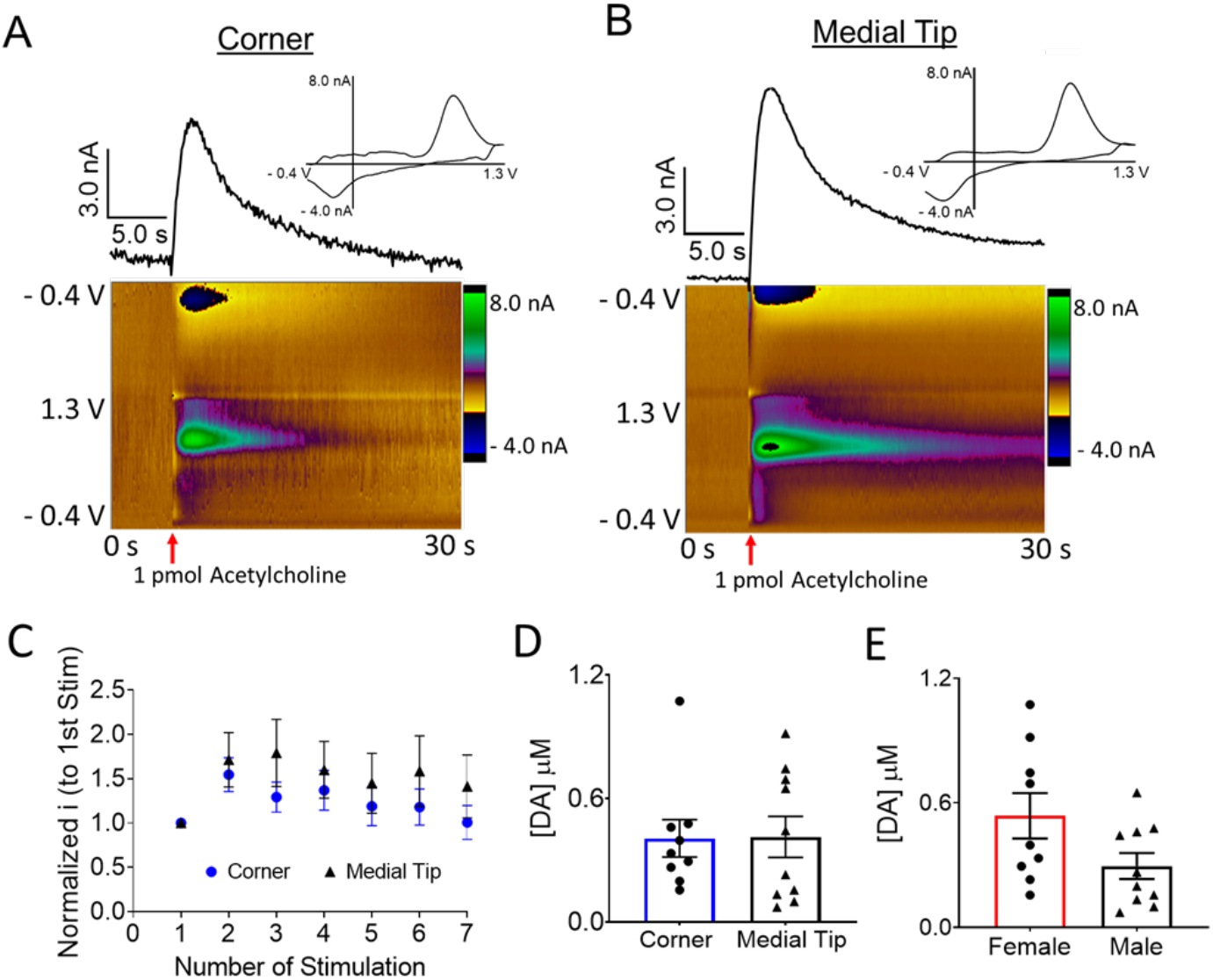
Acetylcholine evoked dopamine release in the MB. Representative FSCV data of stimulated dopamine release in the (A) corner or (B) medial tip of DAT-Gal4, UAS-GFP fly brain. Color plot (bottom) and current vs. time trace (top) show dopamine rises instantaneously upon acetylcholine stimulation (red arrow). The CV (inset) confirms the measured signal is dopamine. (C) Acetylcholine was applied to the corner and medial tip every 10 min to investigate the stability of acetylcholine-evoked dopamine release. Two-way ANOVA shows a significant effect of stimulation number (F _(6,108)_ =5.474, p<0.0001) but not region (F _(1, 18)_ = 0.7110, p = 0.4102). (D) Peak concentration of dopamine release is not significantly different between compartments (n=9 corner, n=10 medial tip, p=0.8676, t-test). (E) Dopamine peak concentrations were not significantly different among sexes (n=9, p=0.0649, t-test).

To investigate stability, 1 pmol acetylcholine stimulation were repeated every 10 min and normalized to the peak current of the first stimulation (Fig. 2C). Dopamine current slightly increased with repeated stimulations at first, an effect we previously observed with acetylcholine and nicotine stimulation in *Drosophila* larval ventral nerve cord ^23^. Two-way ANOVA shows a significant effect of stimulation number for the repeated measurement (F _(6,108)_ =5.474, p<0.0001) but not region (F _(1, 18)_ = 0.7110, p = 0.4102). For average acetylcholine-evoked dopamine release, there were no significant regional differences in dopamine (Fig. 2D, corner, 0.41 ± 0.09 μM, n=9 4 F/ 5 M; medial tip, 0.38 ± 0.10 μM, n=10 5 F/ 5 M, p=0.8676, t-test). Acetylcholine-stimulated release was larger in the MB, which receives a higher density of dopamine innervation, than previous measurements in the central complex (0.26 μM) ^24^. Comparing males and females, the average peak concentration in females was slightly higher (0.54 ± 0.11 μM, n=9) than in males (0.30 ± 0.06 μM, n=10), but it was not significantly different (t-test, p=0.0649).

### ATP/P2X_2_ Mediated Endogenous Dopamine Release in the Mushroom Body of *Drosophila* Brains

Chemogenetic stimulation is a strategy to specifically insert a channel into neurons that is activated by a chemical stimulant. P2X_2_ was expressed in dopaminergic neurons expressing DAT, and GFP was used to visualize the MB for the electrode placement, similar to acetylcholine experiments. The pipette was filled with 1 mM ATP. ATP is electroactive but its oxidation potential is around 1.4 V, thus ATP is not detected with the dopamine FSCV waveform ^22^. Figure 3 shows FSCV data after 0.5 pmol ATP stimulation applied in the MB corner (Fig. 3A) or medial tip (Fig. 3B). As a control, ATP was pressure ejected into the MB of flies not expressing P2X_2_ channels, and no dopamine release was observed (Fig. S1A). Since the coding sequence for P2X_2_ channel was inserted near the DAT, we compared a half-decay time (t_50_) of acetylcholine-evoked dopamine release to explore possible functional alteration on DAT activity and found no significant difference in P2X_2_ flies versus control flies (Fig. S2B, p=0.2034, t-test). Thus, DAT activity did not alter upon genetic modification.

**Figure 3.**
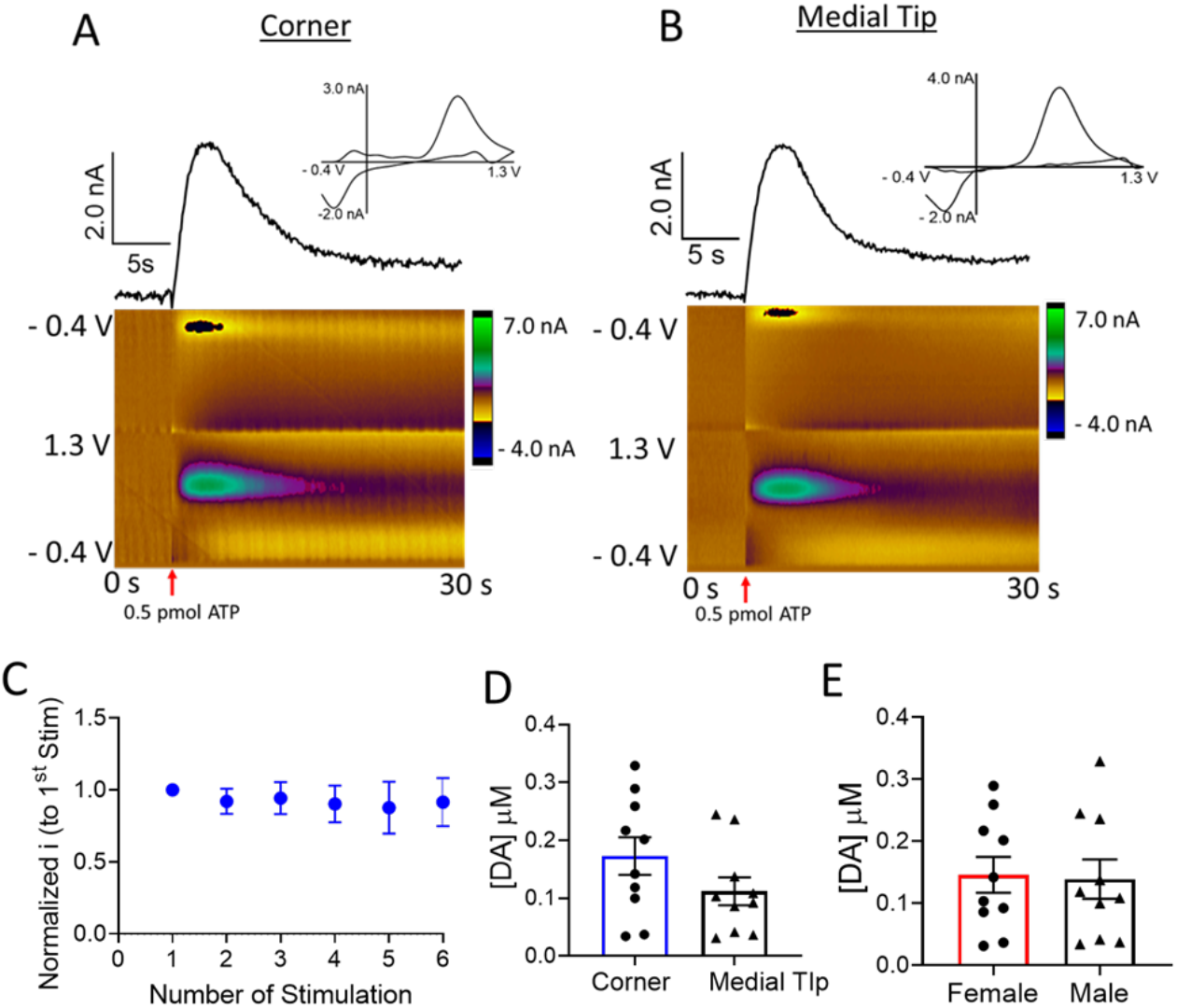
P2X_2_-mediated dopamine release in the MB of UAS-P2X_2_;DAT-Gal4, GFP flies. 0.5 pmol ATP was pressure ejected (red arrow) into the (A) corner and (B) medial tip to elicit endogenous dopamine release. Color plots (bottom), current vs. time traces (top), and CVs (insert) confirm that the measured responses upon ATP stimulations are dopamine. (C) ATP was applied at 10 min intervals to investigate the stability of P2X_2_ stimulated dopamine in the corner. There was no significant effect of stimulation number on dopamine release (F _(5,20)_ = 0.2265, p = 0.9467, One-way ANOVA, n=6). (D) The peak concentration of dopamine release in the corner is not significantly different than that in the medial tip (p=0.1486, n=10 each, t-test). (E) There are no significant differences in peak dopamine concentration by sex (p=0.8711, n=10 each, t-test).

Stability of P2X_2_ mediated dopamine release was investigated in the corner, and release was stable with no effect of stimulation number (Fig. 3C, F _(5,20)_ = 0.2265, p = 0.9467, One-way ANOVA, n=6). P2X_2_ evoked 0.17 ± 0.03 μM dopamine in the corner and 0.11 ± 0.02 μM in the medial tip, which are not significantly different (Fig. 3D, p=0.1486, t-test, n=10). In addition, no significant sex differences were observed (Fig. 3E, female, 0.15 ± 0.03 μM; male, 0.14 ± 0.03 μM, p=0.8711, t-test, n=10).

### CsChrimson Mediated Endogenous Dopamine Release in the Mushroom Body of *Drosophila* Brains

For optogenetic stimulation, light sensitive channels are genetically encoded and action potentials are induced upon photoactivation ^31^. CsChrimson has an excitation maximum in the red, and has been previously been expressed in *Drosophila* larvae for stimulation of endogenous dopamine or octopamine in the ventral nerve cord ^21^. Here, we used optogenetics to study dopamine release and clearance for the first-time *ex vivo* in adult *Drosophila* brains. CsChrimson was expressed in dopaminergic neurons with DAT driver, and red-light induced dopamine was measured in different compartments of the MB with FSCV.

First, a train with multiple pulses of red-light stimulation (20 pulses, 2ms, and 60 Hz) was delivered to elicit dopamine release in the MB corner and medial tip (Fig. 4A and 4B). Dopamine oxidation and reduction are clearly observed in color plots and CVs. Dopamine immediately increases upon the light stimulation and decreases when the light train is finished. As a control, red-light stimulation pulses were delivered to the MB of flies not expressing CsChrimson channels and no dopamine release was observed (Fig. S1B). Next, we investigated a single pulse release by delivering one, 4 ms, 617 nm pulse of light to evoke dopamine in the corner (Fig. 4C) and medial tip (Fig. 4D). The magnitude is about 30 % of the dopamine evoked by 20 pulses, similar to a previous study in the larval VNC where the first light pulse stimulation elicited 20 % of the maximal signal ^21^. The changes in dopamine are very rapid with single-pulse optogenetic stimulation, but the clearance in the corner appears slower than that in the medial tip.

**Figure 4.**
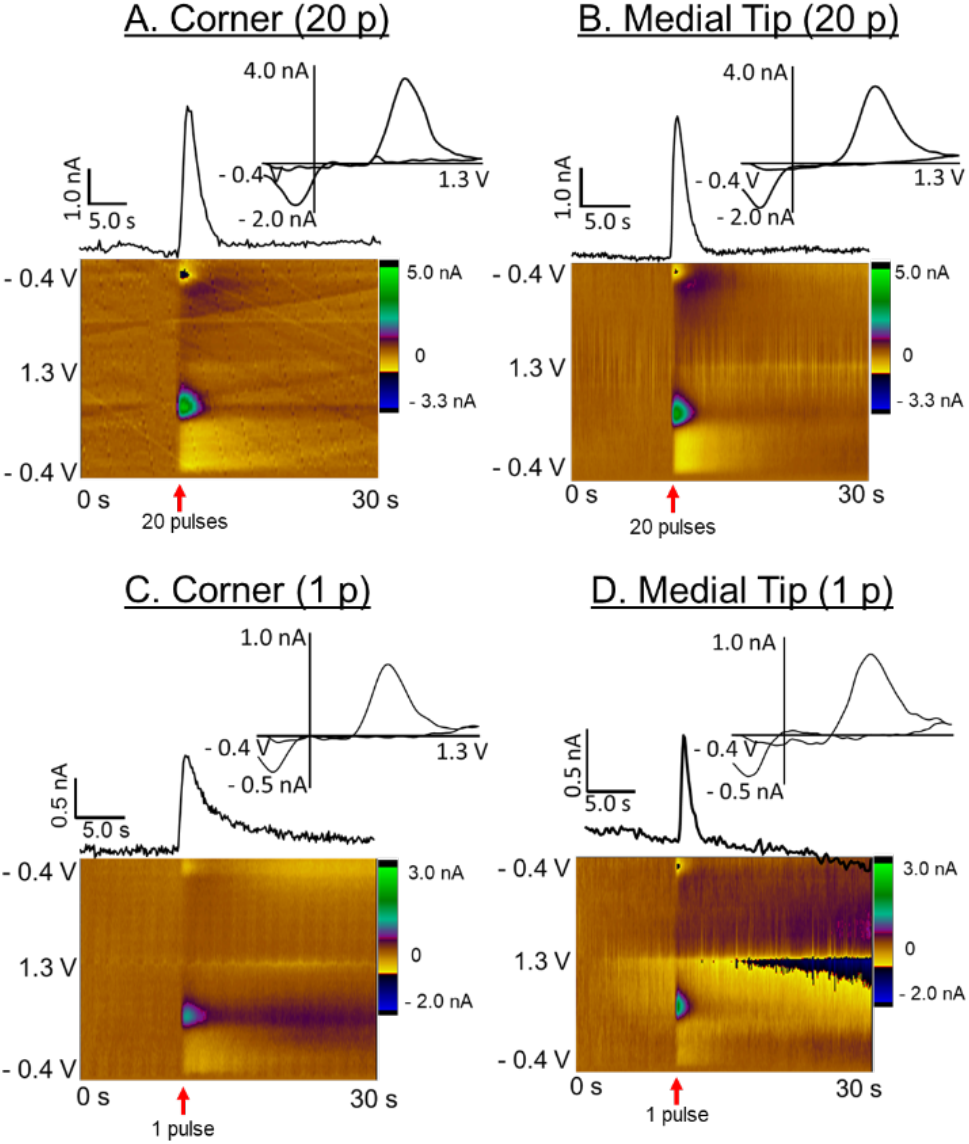
Representative FSCV data of CsChrimson mediated dopamine release upon red-light stimulation. A train of multiple pulses of red-light stimulation (617 nm light, 20 pulses, 2ms, and 60 Hz) was applied (red arrow) to the (A) corner and (B) medial tip in the MB of adult flies expressing CsChrimson in dopaminergic neurons (UAS-CsChrimson; DAT-Gal4, GFP/UAS-Chrismon). A 4 ms single red-light pulse was applied to the (C) corner and (D) medial tip in the MB. All color plots and CVs verify dopamine detection. Current vs. time traces of a single pulse stimulation show slower dopamine clearance in the corner and faster clearance in the medial tip.

Next, we tested the effect of various stimulation parameters on dopamine release in the MB corner. Stimulating pulse width was varied from 1 to 4 ms (20 pulse, 60 Hz). Fig. S3A shows example raw traces for different pulse widths, and the average data in Fig. 5A shows a significant effect of stimulation width on evoked dopamine (F _(3,15)_ =5.844, p=0.0075, n=6, Oneway ANOVA). The average dopamine concentration almost doubled when pulse width was changed from 1 to 2 ms, and 3 and 4 ms also elicited significantly higher dopamine than the 1 ms pulses. Next, the number of stimulation pulses was varied applying 1, 5, 10, 20, 40, 60, and 120 pulses (2ms, 60 Hz, example traces available in Fig. S3B). There was a main effect of pulse number on evoked dopamine release (Fig. 5B, F _(6,30)_ = 6.304, p=0.0002, n=6, One-way ANOVA). The dopamine concentration significantly increased up to 20 pulses then plateaued. The stimulation frequency was varied from 10 to 60 Hz (Fig. 5C, 2ms, 5 pulses). A One-way ANOVA indicates a main effect of pulse frequency (F _(3,12)_ = 8.104, p=0.0034, n=5) and 60 Hz evoked significantly higher dopamine than other frequencies. The stability of CsChrimson mediated dopamine was explored by applying 20 pulses (2 ms, 60 Hz) every 5 or 10 min. Release was stable and there was no main effect of stimulation interval (F_(1, 8)_ = 0.3574, p=0.5665, n=5, Two-way ANOVA) or stimulation number (F_(6,48)_=0.6274, p=0.7075, n=5) on dopamine release.

**Figure 5.**
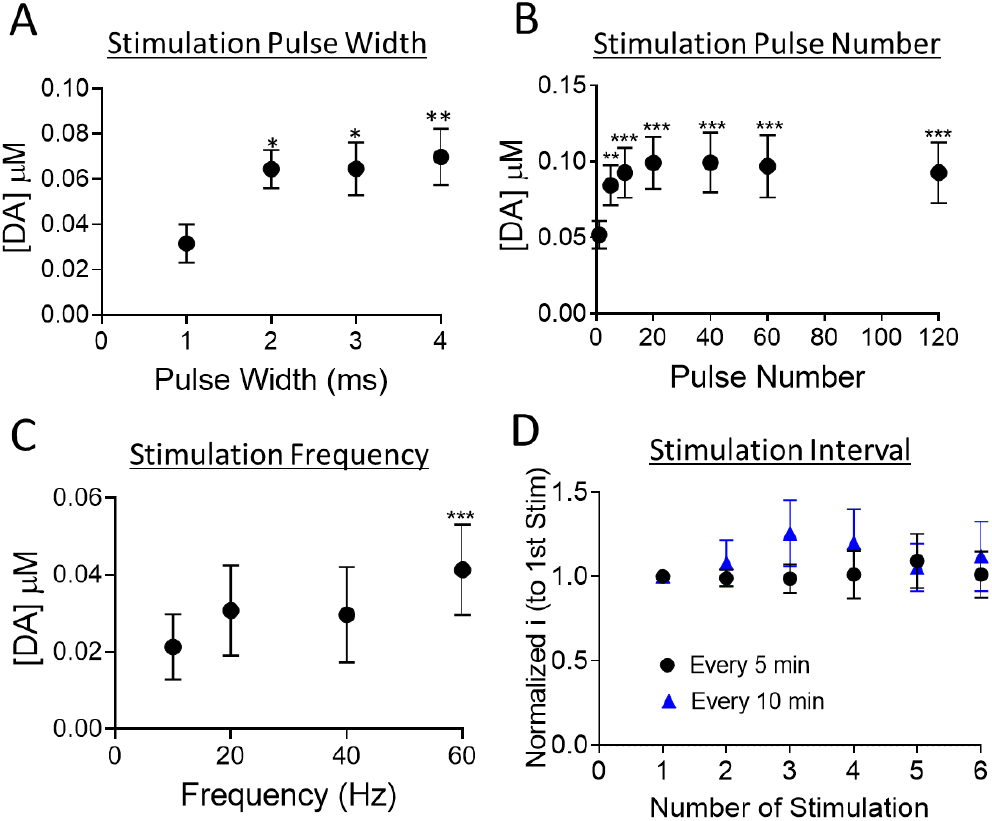
Effect of red-light stimulating parameters on CsChrimson mediated dopamine release in the MB (A) Varying pulse width (20 pulses, 60 Hz). Dopamine peak concentration increased up to 2 ms and then plateaued. There was a significant effect of pulse width (One-way ANOVA, F _(3,15)_ =5.844, p=0.0075, n=6, post-test differences from 1 ms are marked). (B) Varying stimulation pulse number (2 ms width, 60 Hz). Dopamine concentration increased up to 20 pulses and then plateaued. There is a significant effect of pulse number on dopamine release (One-way ANOVA, F _(6,30)_ = 6.304, p=0.0002, n=6, post-test differences from 1 pulse are marked). (C) Varying stimulation frequency (5 pulses, 2 ms). There is a main effect of stimulation frequency (One-way ANOVA, F _(3,12)_ = 8.104, p=0.0034, n=5, post-test differences from 10 Hz are marked). (D) Stability. Optogenetic stimulations (20 pulses, 2 ms, 60 Hz) were applied at 5- or 10-min intervals. Oxidation current is normalized to the first stimulation for each experiment. Release is stable as there is no main effect of stimulation interval (Two-way ANOVA, F _(1, 8)_ = 0.3574, p=0.5665, n=5 each) or stimulation numbers (F _(6,48)_ =0.6274, p=0.7075). * p<0.05, **p<0.01, ***p<0.001

A short train of red-light pulses (20 pulses, 2ms, 60 Hz) was applied to determine regional or sex differences on dopamine release. Optogenetic stimulation evoked on average 0.09 ± 0.01 μM (n=11, 6 F / 5 M) dopamine in the corner, significantly higher than the evoked dopamine in the medial tip (Fig. 6A, 0.05 ± 0.01 μM, p=0.0023, n=17 9 F/ 8 M, t-test). In the corner, significantly differences in dopamine release were evident between sexes, where females (0.12 ± 0.01 μM, n=6) had significantly greater release than males (0.06 ± 0.01 μM, p=0.0047, n=5, t-test). However, no sex differences were observed in the medial tip (female, 0.06 ± 0.01 μM, n=9; male, 0.04 ± 0.01 μM, n=8, p=0.2331, t-test).

**Figure 6.**
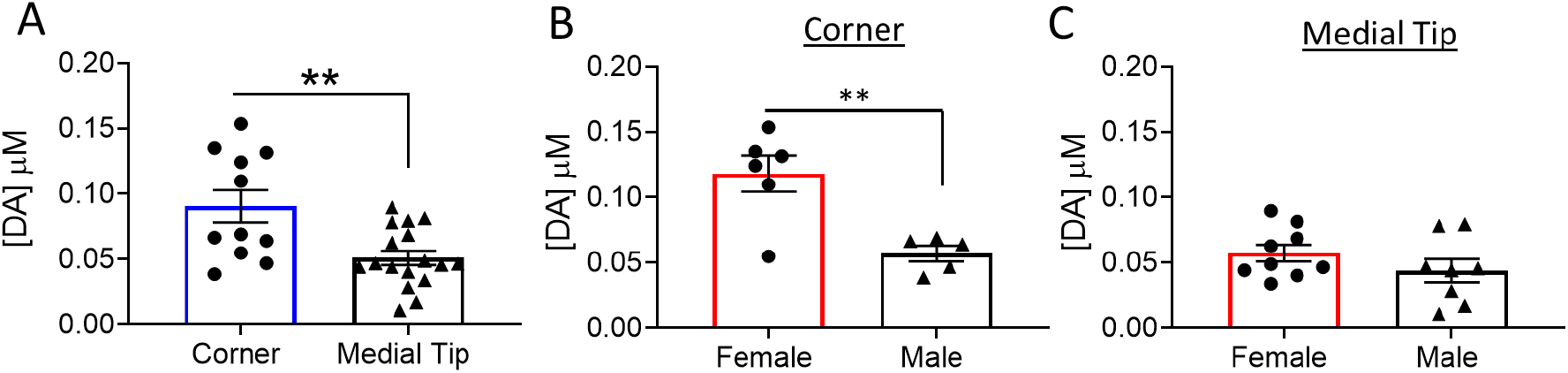
CsChrimson mediated dopamine release in the MB of adult fly brains. Multiple pulses of red-light stimulation (20 pulses, 2ms, 60 Hz) were applied. (A) The peak concentration of dopamine release in the corner is significantly higher than that in medial tip (p=0.0023, corner n=11; medial tip n=17, t-test) (B) Females have significantly higher evoked dopamine release than male flies in the corner (p=0.0047, Female n=6; male n=5, t-test). (C) No significant sex differences in evoked dopamine release were observed in the medial tip (p=0.2331, female n=9; male n=8, t-test)

### CsChrimson stimulation for modeling dopamine uptake in adult Drosophila

One of the benefits of using CsChrimson is that precise pulses are applied and thus we can model the data to obtain the uptake kinetics of released dopamine ^21^. We fit single-pulse evoked dopamine concentration curves to Michaelis-Menten kinetics equation, assuming DAT is the primary mode of clearance, not diffusion (Fig. 7) ^29^. The modeling of single pulse data is advantageous because of the absence of autoreceptor regulation, which might change how much is released per pulse. The model gives a K_m_, a measure of DAT affinity for dopamine, and V_max_, a function of DAT density. There are no significant differences in K_m_ between the corner and the medial tip of MB (Fig. 7A, Corner, 0.23 ± 0.01 μM, n=8 (F 5/ M 3); Medial tip, 0.25 ± 0.01 μM, p=0.684, n=8 (F 4/ M 4), t-test). However, V_max_ is significantly higher in the medial tip (Corner, 0.12 ± 0.01μM/s; Medial tip, 0.19 ± 0.01 μM/s, p<0.05, t-test). The amount of dopamine released per pulse is significantly larger in the corner (Fig. 7C, Corner, 0.05 ± 0.01 μM; Medial tip, 0.03± 0.01 μM, p =0.0056, t-test), similar to the findings of dopamine release with 20 pulse stimulations (Fig. 6B). V_max_ and peak dopamine concentration have an inverse relationship because a faster rate of DAT clearance will lower concentrations of dopamine in the extracellular space. The K_m_ values for dopamine uptake in the adult fly brains are similar to that in the caudate putamen and nucleus accumbens (NAc) of rat brain (0.2 μM) ^32^. In addition, V_max_ in adult flies is the same order of magnitude as previous measurements in larvae ^21,33^.

**Figure 7.**
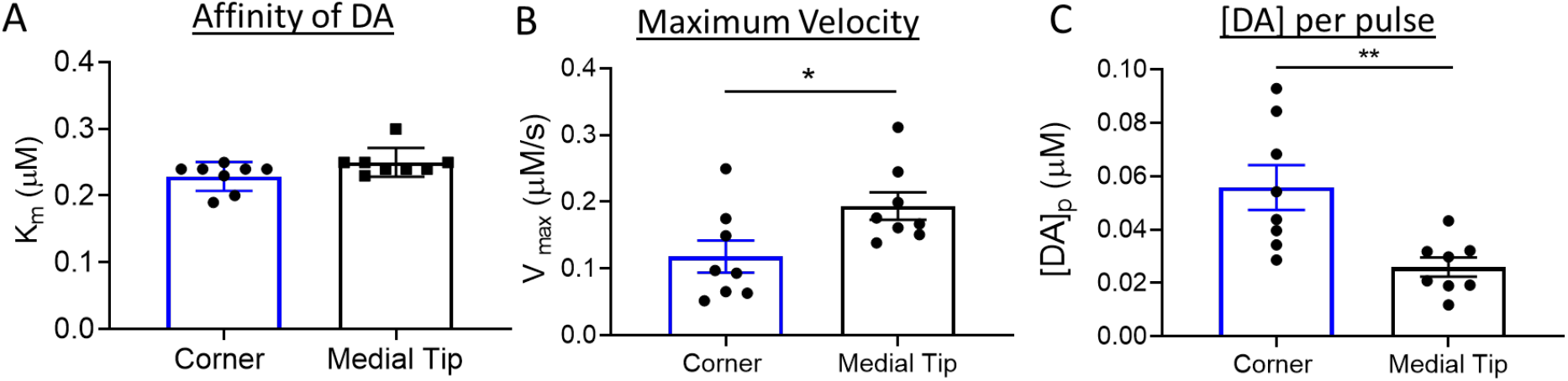
Michaelis-Menten modeling of single-pulse stimulated dopamine release in the MB of adult *Drosophila* brains (Corner, female n=5, male n=3; Medial tip, female n=4, male n=4). (A) No significant difference in affinity of DAT for dopamine (K_m_) was observed between corner and medial tip of MB (p=0.684, t-test) (B) Maximum rate constant for uptake (V_max_) is used to approximate functional DAT density. A significant difference in V_max_ was observed between the corner and the medial tip (p<0.05, t-test) (C) Dopamine per pulse ([DA]_p_) is significantly higher in the corner than that in the medial tip (p =0.0056, t-test).

### Challenges with optogenetic measurements

The biggest challenge for conducting optogenetic experiments with CsChrimson is that the channel is also sensitive to lower wavelengths of visible light ^34^. Compared to Channelrhodopsin-2 (ChR2), CsChrimson’s excitation spectrum is far red shifted, with a maximum excitation at 625 nm, but it also absorbs well at lower wavelengths ^34^. Blue (470 nm) light can still trigger action potentials in the motor neurons in *Drosophila* larvae and stereotyped behavior and adult expressing CsChrimson ^34^.

Electrode implantation is normally facilitated by expressing GFP that is illuminated with blue light (470 nm) to visualize the target area. However, the blue light used to activate GFP causes strong dopamine release by opening CsChrimson, depleting the dopamine pools. Figure 8 demonstrates this effect; current immediately increased and then plateaued when the blue light was turned on for 5 s (blue line). The signal is identified as dopamine by its CV (Fig. 8B). After 30 min rest in the dark, a 20-pulse red-light stimulation was delivered to the same brain, but dopamine release was not observed (Fig. 8A and 8B, red line). White light is typically used during the brain tissue dissection, and it also activates CsChrimson causing dopamine release. Figure 8C shows current response (black line) upon 5 s continuous white light stimulation, and its signal is verified as dopamine (Fig. 8D). However, after 30 min in the dark, a 20-pulse red light stimulation elicits dopamine release (red line), demonstrating that the white light did not fully deplete the dopamine. Practically, we found that it was not possible to place the electrode by using GFP. Instead, we placed it using as low intensity and short duration of white light as possible, and then verified the position of the electrode using GFP at the end of the experiment. Extreme caution must be taken when conducting CsChrimson optogenetic experiments to minimize light exposure of harvested brains. These experiments were very challenging, requiring a highly trained investigator.

**Figure 8.**
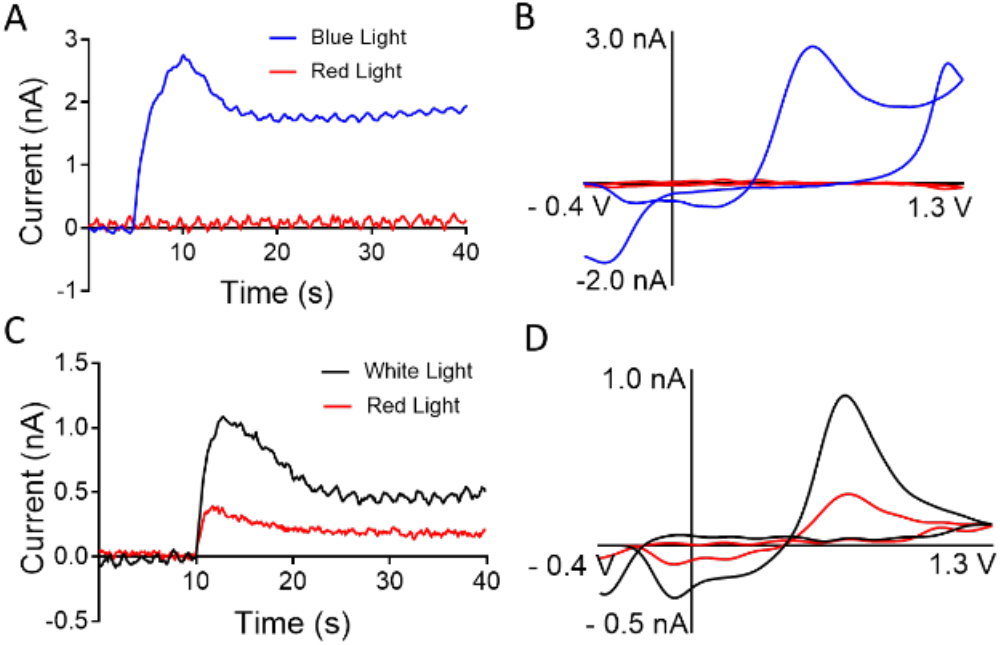
Effect of various colors of light on dopamine release in the MB of adult *Drosophila* brains. (A) Current vs. time trace of light-evoked dopamine release collected upon with 5 s continuous blue light exposure (blue line). In the same subject, 30 min later, no dopamine was observed with multiple pulses of red-light stimulation (red line, 20 pulses, 2 ms, and 60 Hz). (B) Cyclic voltammogram of dopamine release using 470 nm light (blue line) and red-light stimulation (red line). (C) Current vs. time trace of dopamine release upon 5 s continuous white light (black line) and 30 min later (red line) by a train of red-light stimulation pulses (20 pulses, 2 ms, and 60 Hz). (D) Cyclic voltammogram of dopamine release using white light (black line) and red stimulating light (red line, 30 min later).

### Comparison of different stimulation techniques for dopamine measurements in Drosophila adult brains

Different stimulation techniques elicit different concentrations of dopamine in isolated adult brains (Table 1). Acetylcholine stimulation evoked the highest dopamine release, followed by P2X_2_, and CsChrimson. Similar results were obtained in *Drosophila* larval CNS where acetylcholine evoked 0.43 μM dopamine, P2X_2_ 0.4 μM, and CsChrimson 0.15 μM ^21,22,28^. Acetylcholine is a natural neurotransmitter in the brain acting on endogenous presynaptic nicotinic acetylcholine receptors (nAChRs) whereas P2X_2_ or CsChrimson are genetically inserted into the genome and require genetically-altered flies. The number of exogenous P2X_2_ or CsChrimson channels expressed is not known; thus, there may be more nAChRs that modulate dopaminergic neuron activity than P2X_2_ or CsChrimson channels. Because acetylcholine stimulation does not require a genetic manipulation, it is an easy stimulation option to monitor dopamine changes in flies that are genetically modified to model diseases.

**Table 1.**
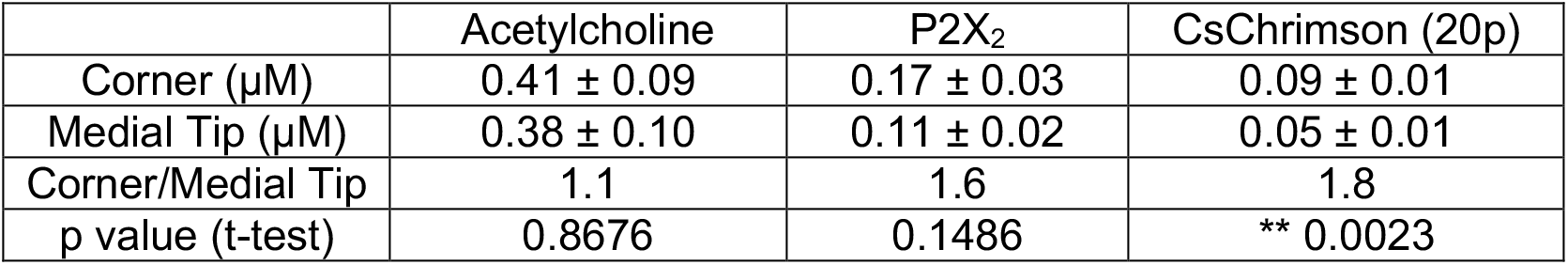
Endogenous dopamine release in the corner and medial tip of mushroom body with different stimulation methods.

One of biggest benefits of using *Drosophila* as model system is the availability of advanced genetic tools to understand the effects of specific genes on physiological functions. Acetylcholine stimulation is non-specific, but P2X_2_ and CsChrimson are genetically expressed specifically only in dopaminergic neurons. Both channels were expressed in dopaminergic neurons with the same DAT driver, and P2X_2_ evoked greater dopamine than CsChrimson. P2X_2_ has a larger channel conductance and is activated a bolus injection of ATP, which stays in the extracellular space longer before it diffuses away or is metabolized ^34,35^. Consequently, large P2X_2_ stimulation could be advantageous for future experiments utilizing more restricted genetic drivers, such as MB compartment or dopaminergic projection pathway specific drivers, that would allow stimulation of discrete sets of dopamine neurons ^4^.

CsChrimson is the only stimulation method that mimics the natural firing patterns of neurons. In mammals, dopaminergic neurons spontaneously fire bursts at higher frequencies (>10 Hz), called phasic firing, and lower frequencies (<5 Hz), called tonic firing, that produce a basal dopamine tone ^36,37^. Here, the stimulation patterns mimic different patterns of phasic firing, but future studies could examine lower frequencies associated with tonic firing. CsChrimson is activated by millisecond-long light pulses to rapidly turn on and off channels, eliciting small amounts of dopamine release that might be more physiological. In addition, the traces resemble those of electrically-stimulated release, which are easier to model uptake parameters. Subtle change in dopamine release were detected with CsChrimson, such as heterogeneity of dopamine regulation in various regions. However, CsChrimson experiments are challenging to perform since CsChrimson is sensitive to ambient white light and blue light used to stimulate GFP fluorescence. Therefore, minimizing the light exposure of isolated brains is essential. Future experiments with an intact, *in vivo* preparation in adult flies might be helpful, where dopamine synthesis would continue to occur after light exposure to reduce depletion. Overall, CsChrimson accomplishes the best spatial and temporal control of neuronal activity, but acetylcholine and P2X_2_ stimulations are easier to implement to monitor changes due to pharmacological agents or between fly lines ^38,22,28^.

### Comparison of sex differences in evoked dopamine release

Table 2 shows a comparison of sex differences in evoked dopamine release in the MB (combined corner and medial tip) with acetylcholine, P2X_2_, or CsChrimson stimulations. The average dopamine concentration in females is higher than males, but significant effects were not observed with acetylcholine or P2X_2_ stimulations. CsChrimson evoked significantly higher dopamine release in female flies, particularly in the corner. CsChrimson stimulations elicit less dopamine release and shorter events, so sex differences in dopamine release may be easier to observe when stimulations are not maximal. These results in *Drosophila* are similar to those for electrically stimulated release in rats, where female rats have higher dopamine tissue content and stimulated release than male rats ^39–41^. In adult *Drosophila*, sex differences in dopamine tissue content have been reported with significant higher dopamine content in females ^42^. Thus, stimulated release is correlated with tissue content, and subtle sex differences are most easily observed with the more sensitive optogenetic stimulations.

**Table 2.** Stimulated dopamine release in the MB of female and male flies with different stimulation techniques.

|  | Acetylcholine | P2X<sub>2</sub> | CsChrimson |
| --- | --- | --- | --- |
| Female (μM) | 0.54 ± 0.11 | 0.15 ± 0.03 | 0.08 ± 0.01 |
| Male (μM) | 0.30 ± 0.06 | 0.14 ± 0.03 | 0.05 ± 0.01 |
| Female/Male DA ratio | 1.8 | 1.1 | 1.6 |
| p value (t-test) | 0.0649 | 0.8711 | * 0.0321 |

### Compartmental differences in dopamine in the mushroom body

The *Drosophila* MB receives projections from different dopaminergic clusters to anatomically and functionally discrete compartments, whose output neurons direct different behaviors ^43^. For example, the corner receives projections from PPL1 (projecting to γ1 and γ2) that control aversive olfactory learning,^6,44^ and the medial tip receives projections from the PAM (projecting to γ3, γ4, and γ5) that control appetitive learning.^7,45^. Here, subtle differences were observed between mushroom body compartments with optogenetic stimulation. Larger dopamine signaling was observed in the corner, which processes aversive learning, compared to the medial tip, which modulates appetitive learning. Thus, aversive learning may require a stronger dopaminergic response. The different compartments also had differences in uptake, which shows differential control of dopamine by DAT between compartments. The corner has slower V_max_ and higher dopamine release compared to the medial tip, while the average K_m_ did not change. The higher V_max_ in the medial tip is likely due to higher expression of DAT and higher uptake serves to dampen dopaminergic signaling. Other studies have linked DAT expression with behavior, for example overexpression of DAT in the MB abolished olfactory aversive memory ^46^. Similarly, we observed lower DAT activity in the MB γ1 and γ2 compartments, which are linked to aversive outputs. Lower dopamine release is generally due to lower concentrations in vesicles or a smaller proportion released per exocytosis event, and the medial tip had lower release as well. Interestingly, the number of dopaminergic neuronal cells that project to the corner is lower than that to the medial tip, suggesting the density of dopaminergic innervation is not correlated with release ^4,47^. Future experiments could also explore autoreceptor regulation or synthesis to understand their contribution to dopaminergic signaling.

To better understand dopaminergic signaling in each MB compartment, future experiments could use MB compartment drivers to express either P2X_2_ or CsChrimson in specific compartments^48,49^. These experiments would precisely measure differences in rapid dopamine signaling, which could better be linked to regulation of specific behaviors. The tools developed here will be useful for future experiments *in vivo* as well, delineating how dopaminergic signaling in discrete compartments regulates behavior as well.

## Conclusions

We employed different stimulation techniques to evoke endogenous dopamine release within different compartments of the mushroom body (MB) for the first time. There were no significant regional or sex differences in evoked dopamine release when acetylcholine and P2X_2_ stimulations were employed but with CsChrimson, dopamine release was significantly higher in the MB corner and in female flies. Acetylcholine stimulated the largest dopamine release, followed by P2X_2_, and CsChrimson stimulation. Acetylcholine stimulation does not require any genetic manipulations so it can performed in any fly line, including disease models. P2X_2_ and CsChrimson use genetic approaches and thus, the channels are expressed in specific neurons. CsChrimson uses offers precise temporal control of neuronal activity but the optogenetic experiments are challenging because CsChrimson is activated at many frequencies of light during dissection and electrode placement. Thus, the light exposure of the isolated fly brains must be minimized. Here, we have developed a toolkit of stimulation approaches, introducing chemogentic and optogenetic experiments for the first time, for measuring endogenous dopamine release in *Drosophila*. These methods demonstrated compartmental differences in MB dopamine and can be used in the future improve our understanding of neurochemical signaling in *Drosophila*.

## Author Information

### Funding

This work is funded by NIHR01MH085159 to BJV.

### Note

The authors declare no competing financial interest.

## Acknowledgments

Authors would like to thank Dr. Charles Nicholson and Dr. Jyoti C. Patel at the New York University School of Medicine for sharing dopamine kinetic modeling MATLAB codes.

## Notes

### Competing Interest Statement

The authors have declared no competing interest.

