## Supporting Information for "Real-Time Measurement of Stimulated Dopamine Release in Compartments of the Adult *Drosophila melanogaster* Mushroom Body"

Mimi Shin, Jeffery M. Copeland, and B. Jill Venton*

Department of Chemistry, University of Virginia, Charlottesville, Virginia 22901

I. Supplemental Methods

II. Supplemental Figures

### I. Supplemental Methods

#### Chemicals

All chemicals were purchased from Sigma-Aldrich (St. Louis, MO) unless otherwise noted. Aqueous solutions were made with Milli-Q water (Milli-pore Billerica, MA). For calibrations, a phosphate buffer (PBS) solution was used (131.25 mM NaCl, 3.0 mM KCl, 10.0 mM $NaH_2PO_4$, 1.2 mM $MgCl_2$, 2.0 mM $Na_2SO_4$, and 1.2 mM $CaCl_2$, pH adjusted to 7.4). A dopamine stock solution was prepared in 0.1 M $HClO_4$ and diluted to 1.0 μM with PBS for calibrations on the day of experiment. For *Drosophila* dissection and FSCV experiments, 11.1 mM glucose and 5.3 mM trehalose were added to PBS prior to each experiment. Stimulation solutions of 5 mM acetylcholine and 1 mM ATP were also prepared in PBS. All-trans-retinal (ATR) stock solution was prepared in ethanol, then kept at – 20 °C, and diluted to 400 μM in pre-made fly food.

#### *Drosophila melanogaster* Strains

DAT-Gal4 (Stock 48359), UAS-CsChrimson (stock 55134), UAS-P2$X_2$ (Stock 76032), UAS-CD8-GFP (stock 5130), were purchased from the Bloomington Drosophila Stock Center (Indiana University, Bloomington, IN). A stable 3$^{rd}$ chromosome DAT-Gal4, UAS-GFP strain was created by crossing heterozygous females to the TM3, Sb balancer chromosome. Progeny carrying both DAT-Gal4 and UAS-GFP elements, a product of meiotic recombination, were checked for stable mushroom body GFP expression and self-crossed to maintain the stock. A stable line carrying two copies of UAS-CsChrimson on separate chromosomes was created by first individually crossing a 2$^{nd}$ chromosome and a 3$^{rd}$ chromosome UAS-CsChrimson element to the CyO/Sp; TM3, Sb/TM6b, Hu double balancer.  Progeny were then crossed to select for the lines carrying balanced UAS-CsChrimson elements.

To generate a *Drosophila* line expressing P2$X_2$ in dopaminergic neurons, DAT-Gal4, GFP was crossed with UAS-P2$X_2$. For optogenetic experiment, DAT-Gal4, GFP line was crossed with a homozygous CsChrimson line. The offspring of optogenetic flies were collected and transferred into a fly vial containing 400 μM all-trans-retinal (ATR) food and flies were fed for 5 days prior to the experiment. ATR is light sensitive; thus, vial is shield from the light. To keep the ATR containing food fresh, the flies were transferred to a new ATR food every two days.

#### Electrochemical measurements

Carbon fiber microelectrodes were fabricated with T-650 carbon fibers (Cytec Engineering Materials, West Patterson, NJ) and prepared as described in Shin *et. al.* [24]. Briefly, a 7 μm carbon fiber was aspirated into a glass capillary (1.2 mm O.D and 0.68 mm I.D, A-M system, inc, Carlsberg, WA) which was pulled into two electrodes using a PE-22 heated coil puller (Narishige int, USA, East Meadow, NT). The carbon fiber was trimmed about 50 μm away from the end of pulled glass tip. A trimmed carbon fiber is sealed with epoxy with Epon Resin 828 (Miller-Stephenson, Danbury, CT) mixed with 14 % (w/w) m-phenylenediamine hardener (Fluka, Milwaukee, WI) and excess epoxy was cleaned with acetone. Sealed electrodes were cured at 100 °C for 2 hours followed by 150 °C overnight. Prior to each experiment, electrodes were soaked in isopropanol for 10 min. Electrical connection was completed by backfilling the electrode with 1M KCl. FSCV was performed using a ChemClamp potentiostat (Dagan, Minneapolis, MN), PCI 6711 and 6052 computer interface cards (National Instrument, Austin, TX), and a home-built breakout box. For the data collection and analysis, TarHeel CV (provided by R.M. Wightman, University of North Carolina) was used. A waveform of – 0.4 V to 1.3 V and back to – 0.4 V at 400 V/s to a CFME for dopamine detection.

#### Imaging of Drosophila Brain Tissues

The procedure of the brain tissue and mounting preparation for imaging was adapted from previously reported literature [50]. Isolated brains were transferred into a 1.5 mL microcentrifuge tube containing 4 % formaldehyde and incubated for 20 min with slow speed rocking at room temperature. Following tissue fixation, the formaldehyde was removed and fixed brain tissues were washed with PBS three times; one, 20 min long wash and two, 5 min quick washes. PBS solution was changed between washes. To mount the brain tissue, a bridge slide was constructed by placing two small cover slips (Ted Pella, Redding, CA) on each side of a glass microscope slide (Fisher Scientific, Hampton, NH) roughly 1 cm apart. Then, the three outer edges of each cover slide were sealed with clear nail polish. Then, the gaps between cover slips were filled with Vectashield mounting media (Vector Lab, Burlingame, CA). Using forceps, individual brain tissue was transferred and positioned anterior slide facing up. Another cover slip was placed over the base cover slip and the brain tissue. The bridge slip was sealed with clear nail polish, and the brains were imaged using a Nikon A1Rsi upright confocal microscope (Nikon Instruments Inc, Melville, NY) with GaAsP detectors and 10X Plan Apo NIR WD objective. Z-stack image was taken with 10 μm increments z steps. The viability of isolated brain was previously demonstrated using calcein imaging [24].

#### II. Supplemental Figures

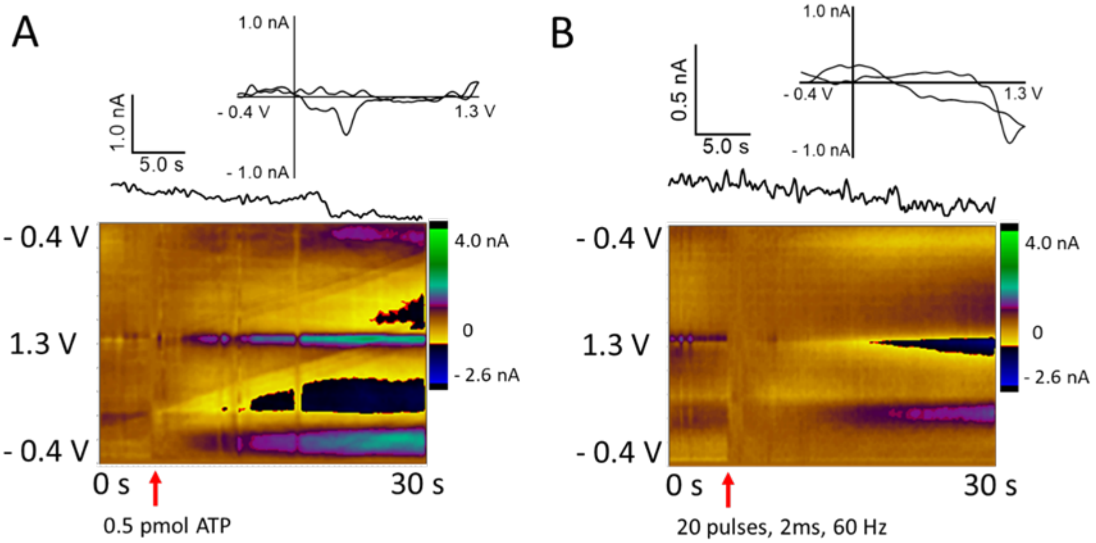


**Supplementary Figure 1.** Control experiments for $P2X_2$ and CsChrimson mediated dopamine release. (A) 0.5 pmol ATP was pressure ejected into the corner of DAT-Gal4, GFP flies. Upon ATP stimulation, no dopamine release was observed. (B) Multiple pulses of red light (20 pulses, 2 ms, and 60 Hz) were applied to the corner of DAT-Gal4, GFP flies. The color plot (bottom), current vs. time traces, and CV identified that no dopamine was released upon the light stimulation.

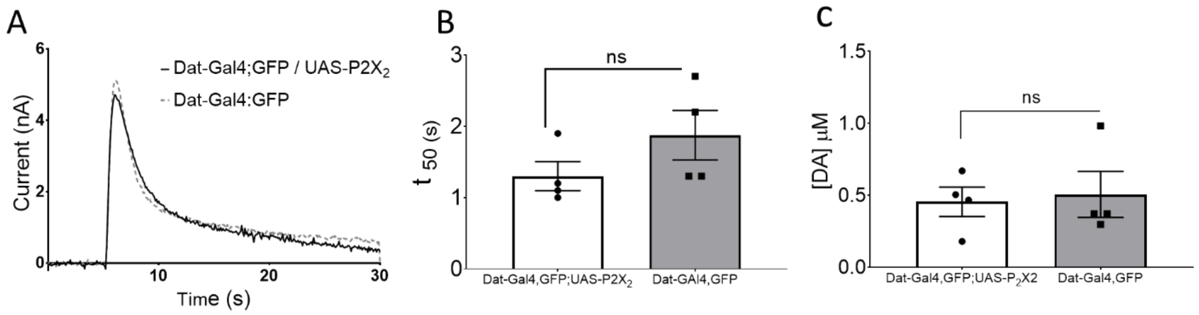


**Supplementary Figure 2.** Characterization of dopamine clearance in DAT-Gal4, GFP;UAS-P2X2 ($P2X_2$) and DAT-Gal4, GFP (Control) flies using 1 pmol acetylcholine stimulations. (A) Current vs. time traces of acetylcholine-stimulated dopamine release in $P2X_2$ and control flies. (B) Dopamine clearance rate can be approximated using the half-decay time ($t_{50}$), the time it takes the response to reach from the maximum to the half of the peak concentration. $t_{50}$ in $P2X_2$ flies is not significantly different than that in control flies ($P2X_2$, 1.3 ±. 0.2 s; control, n=4, 1.8 ± 0.3 s, p=0.203, t-test, n=4 each) (C) The average peak concentration of acetylcholine-stimulated dopamine release in $P2X_2$ is not significantly different than control ($P2X_2$, n=4, 0.46 ± 0.10 μM; control, 0.50 ± 0.16 μM, p=0.7971, t-test).

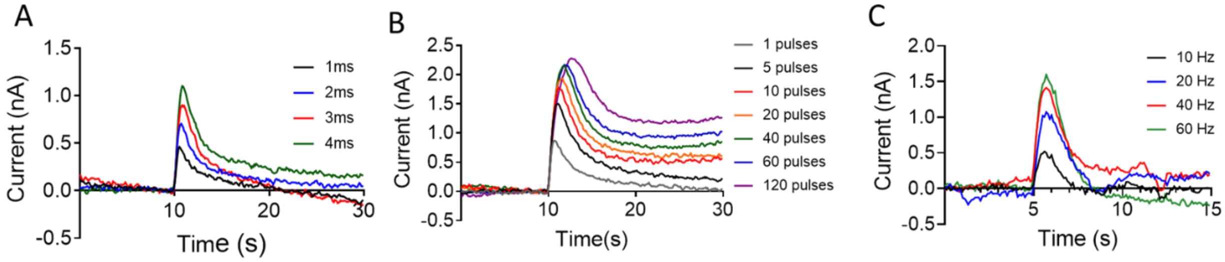


**Supplementary Figure 3.** Current vs. time traces of evoked dopamine release at various (A) stimulating width (1, 2, 3, 4 ms, 20 pulses, 60 Hz), (B) pulse number (60 Hz, 2 ms width), and (C) frequency (5 pulses, 2 ms width).
